# CHAI: Consensus Clustering Through Similarity Matrix Integration for Cell-Type Identification

**DOI:** 10.1101/2024.03.19.585758

**Authors:** Musaddiq K Lodi, Muzammil Lodi, Kezie Osei, Vaishnavi Ranganathan, Priscilla Hwang, Preetam Ghosh

## Abstract

Several methods have been developed to computationally predict cell-types for single cell RNA sequencing (scRNAseq) data. As methods are developed, a common problem for investigators has been identifying the best method they should apply to their specific use-case. To address this challenge, we present CHAI (consensus Clustering tHrough similArIty matrix integratIon for single cell type identification), a wisdom of crowds approach for scRNAseq clustering. CHAI presents two competing methods which aggregate the clustering results from seven state of the art clustering methods: CHAI-AvgSim and CHAI-SNF. Both methods demonstrate improved performance on a diverse selection of benchmarking datasets, besides also outperforming a previous consensus clustering method. We demonstrate CHAI’s practical use case by identifying a leader tumor cell cluster enriched with CDH3. CHAI provides a platform for multiomic integration, and we demonstrate CHAI-SNF to have improved performance when including spatial transcriptomics data. CHAI is intuitive and easily customizable; it provides a way for users to add their own clustering methods to the pipeline, or down-select just the ones they want to use for the clustering aggregation. CHAI is available as an open source R package on GitHub: https://github.com/lodimk2/chai

## 1 Introduction

The advent of single cell RNA sequencing (scRNAseq) has allowed researchers to investigate transcriptional mechanisms at the single cell resolution. Notably, scRNAseq has contributed to the identification of rare cell types, assessing cell heterogeneity, and quantifying cell-cell variation [1]. A common methodology for identifying subpopulations from single cells has been unsupervised clustering [2]. However, the nature of scRNAseq data presents unique challenges in identifying accurate clusters. For example, scRNAseq data is sparse, with frequent gene and cell dropouts. Additionally, scRNAseq data is high dimensional, which tends to data points being similar and therefore unreliable for downstream clustering tasks. Due to these factors, a diverse array of scRNAseq clustering methods have emerged recently [2].

While several clustering methods for scRNAseq data have been published, comprehensive benchmarking studies, such as the one from Yu et al., have indicated that there is no clear “best method” across all scenarios [3]. Due to the high amount of variability in scRNAseq data, even the most commonly used clustering algorithms have distinct strengths and weaknesses. Take for example Seurat, perhaps the most commonly used scRNAseq clustering platform: while results from Seurat often demonstrate high concordance with ground-truth cell type populations, it also tends to overestimate the number of distinct cell types in a dataset [3] [4]. Seurat, along with other popular scRNAseq clustering workflows such as Spectrum and SC3, use community detection algorithms such as Leiden and Louvain as the primary mechanism for their clustering. Preprocessing steps, such as highly variable gene selection, or dimensionality reduction through Principal Component Analysis (PCA), have also become common place before performing the final clustering [3] [4] [5] [6]. Additionally, common unsupervised clustering algorithms, such as *k* means or hierarchical clustering, are used to create initial clusters before reclustering, such as in CIDR [7]. More recently developed algorithms such as scSHC and CHOIR use a statistical significance testing to determine final cluster assignments, and also serve as an evaluation framework outside of the commonly used metrics such as Adjusted Rand Index (ARI) and Normalized Mutual Information (NMI) [8] [9] [10] [11] [12].

With the various scRNAseq clustering methodologies currently available, a common question for investigators becomes: Which method should I use? As there is no definite answer for this, an intuitive approach is to integrate the results from the different clustering algorithms, into a “clustering ensemble” or “consensus clustering” [13]. This idea extends from the wisdom of crowds approach, which states that knowledge from the collective of a group is greater than that of an individual [14] [15] [16]. The idea of consensus clustering was introduced by Strehl and Ghosh, who pioneered hypergraph partitioning algorithms for integrating results from individual clustering results [17]. A method known as SAFE-Clustering implemented all three of Strehl and Ghosh’s algorithms in an application to scRNAseq clustering, which included the clustering methodologies Seurat, SC3, CIDER, t-SNE + k-means in 2018 [18]. SAFE-Clustering demonstrated robust performance across 12 benchmarking datasets, establishing the premise that consensus clustering is applicable to scRNAseq data. Another ensemble clustering method, SAME-Clustering, uses a Mixture model Ensemble to aggregate results from different scRNAseq clustering methodologies [19]. However, since these methods were created in 2020 and prior, there have been further advancements made to the existing algorithms in their pipeline such as Seurat and SC3, and the other algorithms, such as CIDER and SIMLR, are not as widely used [3]. Additionally, these ensemble clustering approaches are not immediately extendable to multi-omic data integration, which can provide even more insights towards distinct cell types and state. A consensus aggregation approach is only as accurate as the performance of the individual information, and so we identified a need for an updated consensus clustering framework that can also seamlessly allow for multiomic data integration.

Here we present CHAI (consensus Clustering tHrough similArIty matrIces), a consensus clustering methodology built upon binary similarity matrices. CHAI contains two clustering ensemble approaches, named CHAI-AvgSim and CHAI-SNF. CHAI-AvgSim is performed by aggregating all clustering assignments with an average similarity matrix, and performing Spectral Clustering on the final average matrix. CHAI-SNF extends Similarity Network Fusion (i.e., SNF), which is a network integration algorithm originally designed for multiomic data integration for patient subtyping and classification [20]. Both CHAI methods have demonstrated improved performance across several benchmarking datasets and conditions, showcasing limited variability across runs, and low impact from poor performing algorithms. Additionally, we present a technique to integrate other data modalities into the CHAI framework, such as spatial transcriptomic data or ATAC-Seq data. CHAI contains seven state of the art scRNAseq clustering algorithms (Seurat-Louvain, Seurat-SLC, CHOIR, RACEID, SC3, Spectrum, and scSHC), and is available as an R package [4] [8] [21] [5] [6] [9]. We seek to make CHAI a collaborative tool for the community by providing a way for scientists and developers to integrate their own clustering algorithms into the pipeline as well, which may potentially strengthen results as more advanced scRNAseq clustering algorithms emerge in the future.

## 2 Results

### 2.1 CHAI Workflow

CHAI is a consensus clustering method that presents two different approaches for the integration of individual clustering results: Average Similarity and SNF [20]. For a more detailed description of each method, please refer to the Methods section.

All CHAI related methods (CHAI-AvgSim, CHAI-SNF, and CHAI-ST), operate under binary matrices. For clustering algorithms, these matrices are calculated by determining if two cells are predicted to be in the same cluster. If they are, we assign 1 to the matrix entry to designate that these two cells are related. If not, we assign 0.

For the spatial coordinates binary matrix representation, we use the methodology from GraphST [22]. First we calculate a pairwise distance between cells based on the spatial coordinates. Then we run a KNN graph, with K being 3. If two cells are neighbors based on this KNN graph, we assign a value of 1 to this cell-cell relationship. If not, we assign 0.

To further illustrate this concept, consider a toy example with three clustering algorithms, and three cells. For all CHAI methods, we first calculate the binary matrices. Fig. 1a depicts the overall workflow while Figs. 1b and 1c show the example runs of the CHAI methodology.

**Figure 1.**
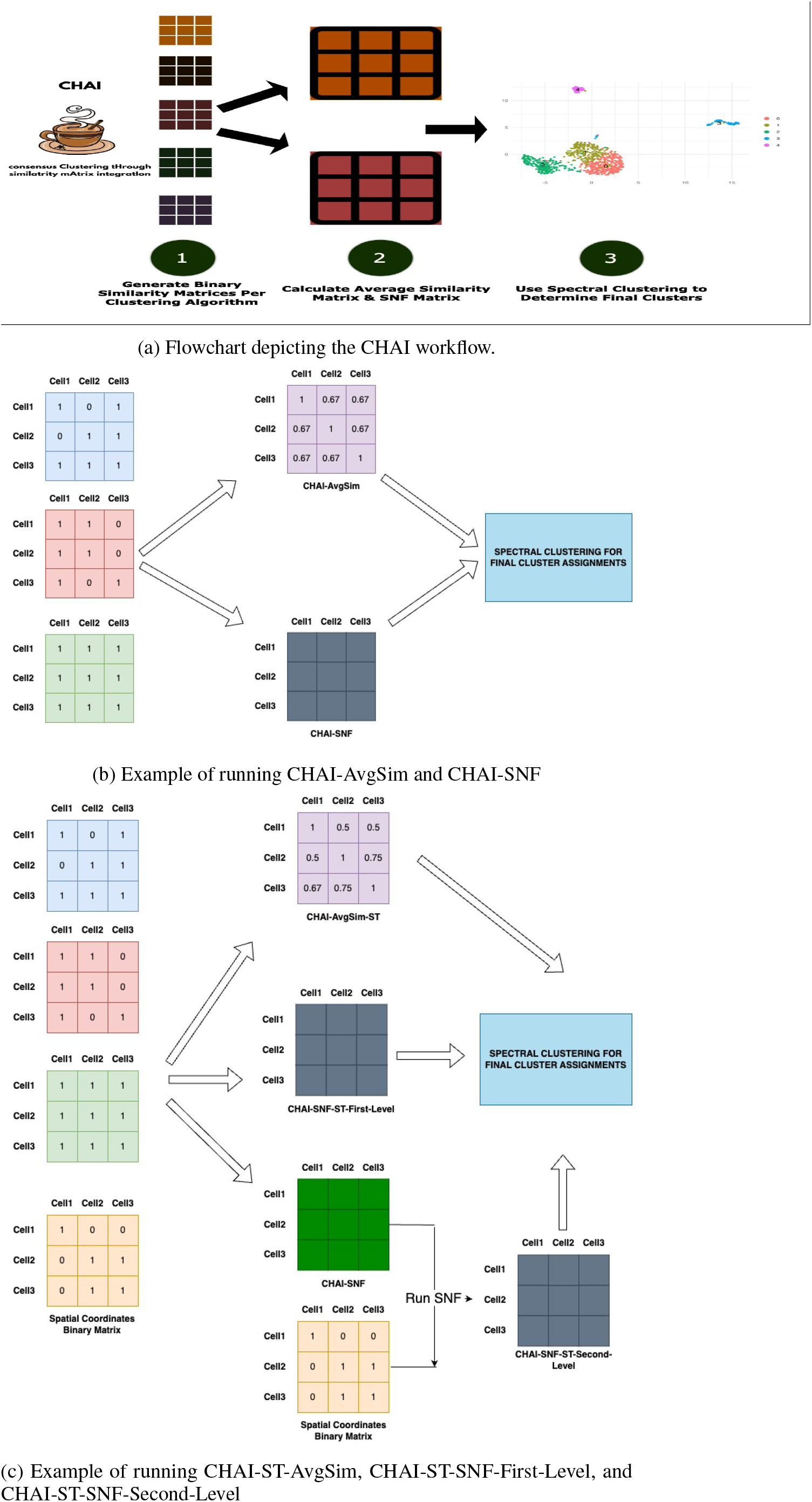
CHAI Workflow Examples

Fig. 1b shows how CHAI-AvgSim and CHAI-SNF are run. For CHAI-AvgSim, we calculate an average of all the binary matrices from the different clustering algorithms. Then, we run Spectral Clustering on the resultant matrix to determine the final clusters. For CHAI-SNF, we run SNF with default parameters on the binary matrices from the clustering algorithms. Then we perform Spectral Clustering on the resultant CHAI-SNF matrix to determine the final clustering assignments.

We present three different ways to integrate spatial transcriptomic data into CHAI (Fig. 1c). For CHAI-AvgSim-ST, we simply include the binary matrix representation of the spatial coordinate data as another matrix to be included into the AvgSim calculation. We then run Spectral Clustering on the resultant matrix to determine the final clusters. For CHAI-SNF-First-Level, we run SNF on all binary matrices, including the spatial coordinates binary matrix representation. For CHAI-SNF-Second-Level, we first run SNF on just the clustering assignment matrices, and keep the spatial coordinates binary matrix separate. Once the SNF matrix for the clustering assignment binary matrices are calculated, we run SNF again, this time with the clustering assignment matrix from the first level SNF and the spatial coordinates matrix. For both CHAI-SNF-First-Level and CHAI-SNF-Second-Level, we run Spectral Clustering on the resulting matrix to determine the final clusters. The main difference between CHAI-SNF-First-Level and CHAI-SNF-Second-Level is that the latter gives more weight to the spatial coordinate data, since it is included separately as an “omic” rather than just another clustering assignment as considered in CHAI-SNF-First-Level. Users may make the decision to run CHAI-SNF-First-Level or CHAI-SNF-Second-Level based on their prior biological knowledge of their datasets.

We benchmarked the performance of both CHAI methods on several datasets. We used 10 publicly available scRNAseq datasets for our main performance evaluation. Additionally, we took advantage of the size and complexity in the Zheng68K PBMC dataset to create subsampled datasets to evaluate the performance of CHAI on various dataset conditions, such as number of cells and number of cell types. In brief, we find that CHAI is a more consistent and accurate performer in diverse dataset conditions when compared to baseline algorithms.

We chose to evaluate using ARI and NMI as they each measure the overlap between predicted and ground truth clustering assignments, and their value decreases as disagreements between sub-populations increase [23]. We display the ARI evaluation in the main text, and the NMI evaluation in the supplementary materials.

### 2.2 CHAI outperforms existing clustering methods on benchmarking datasets

To assess the performance of CHAI-AvgSim and CHAI-SNF, we compared them to seven individual algorithms that form the consensus method. We ran each algorithm on 10 commonly used benchmarking datasets with varying tissue source, number of cells and number of cell types. We evaluated the performance using Adjusted Rand Index (ARI).

We see in Fig. 2a that both CHAI-AvgSim and CHAI-SNF demonstrate robust and consistent performance across benchmarked datasets. Notably, CHAI-AvgSim was a top three performer in 8 out of 10 datasets. We show the frequency of top three performers in each dataset in a heatmap, depicted in Fig. 2b. CHAI-AvgSim and CHAI-SNF have the highest frequency of being the top three performing algorithms, with scores of 80% and 60% respectively.

**Figure 2.**
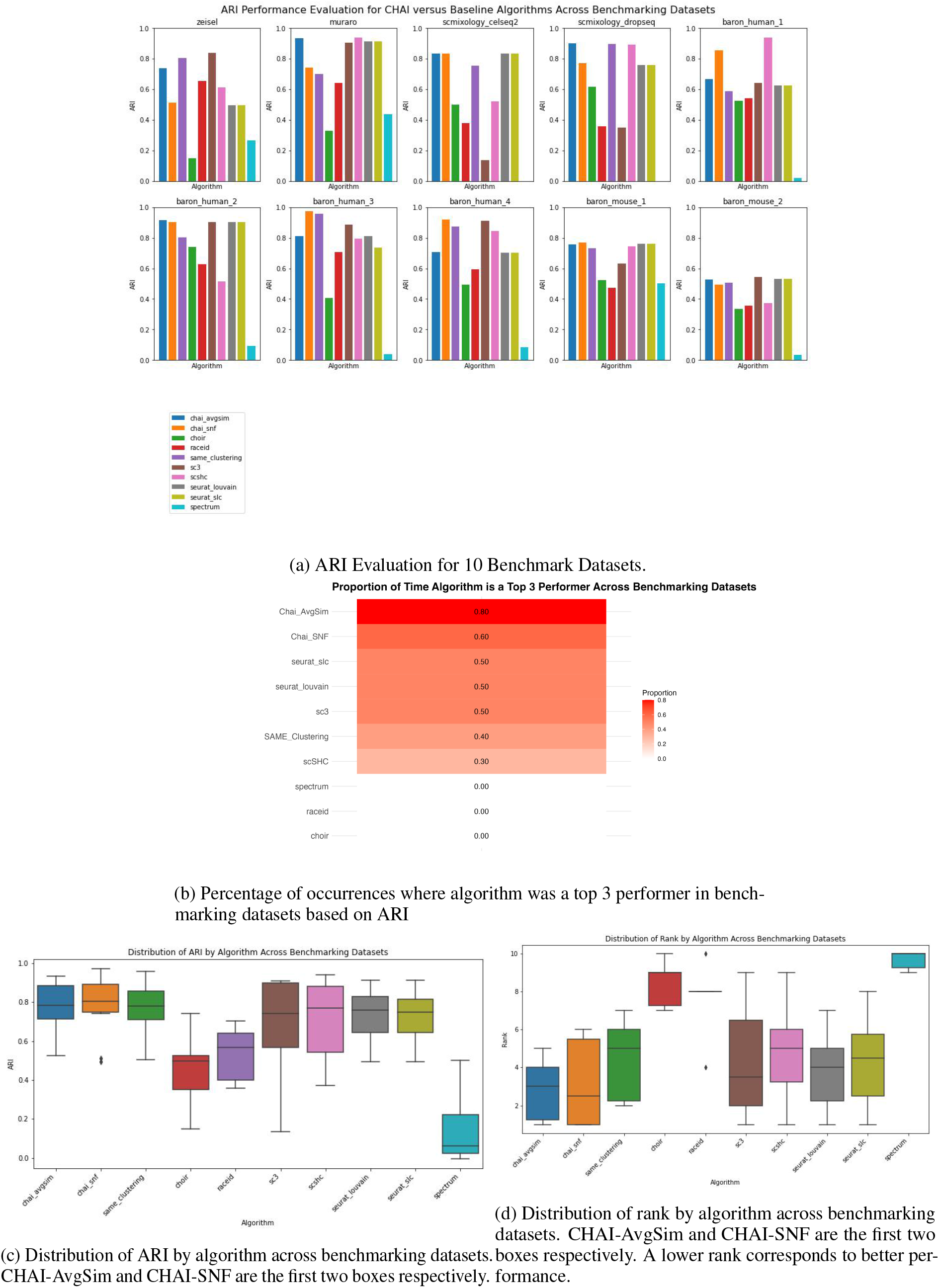
CHAI Evaluation on Benchmarking Datasets

The variability of performance in other baseline algorithms is very noticeable in this analysis. Widely used algorithms such as SC3 and RaceID demonstrate very strong performances in some datasets, like the Zeisel mouse brain dataset, but very poorly in others, such as the SC-Mixology-Dropseq dataset [21] [5] [24] [25]. The primary benefit of the CHAI consensus algorithms is that they reduce this variability in performance. We visualize this variability by plotting the distribution of ARI values as a boxplot, seen in Fig. 2c. Both CHAI-AvgSim and CHAI-SNF have higher median ARI than any of the baseline clustering methods. This analysis also helps to highlight the difference in performance between the two CHAI methods. CHAI-SNF has a higher median ARI, a higher third quartile threshold, and a higher maximum ARI than CHAI-AvgSim, demonstrating its potential for high accuracy. However, it has a much larger interquartile range, which suggests higher variability in performance. CHAI-AvgSim, on the other hand, has a comparable median ARI with other baseline methods, such as Seurat-Louvain and Seurat-SLC. The primary advantage of CHAI-AvgSim lies in its low interquartile range, as it has the lowest interquartile range when compared to any other baseline algorithm. This shows that CHAI-AvgSim is a much more consistent performer across various datasets than any other algorithm including CHAI-SNF, making it a robust choice.

We also calculated the rank of each algorithm across the benchmarking datasets, as shown in Fig. 2. This was done as another metric to measure top performance. Algorithms with a lower rank are higher performers (1 being the best rank, and so on). The median rank of CHAI-AvgSim and CHAI-SNF are quite low at approximately 3 making them a safe choice for accurate clustering across diverse datasets. Additionally we see that the minimum for CHAI-SNF and CHAI-AvgSim is approximately 1 and 2 respectively, showing that it is more likely to be a top performing algorithm than the other baseline algorithms.

Here we also compare CHAI to a previous consensus clustering method, SAME Clustering [19]. CHAI incorporates more algorithms than SAME clustering, and also runs the latest version of Seurat [4]. We demonstrate that at least one of the two CHAI methods outperforms SAME clustering in 8 of the 10 datasets. SAME clustering and CHAI have similar median ARI’s and distributions. CHAI-SNF has the highest upper quartile cutoff value, and the highest median across all algorithms. It also demonstrated the highest ARI for any of the benchmarking datasets. Despite the similarities in ARI distribution, we see that both CHAI methods have a lower distribution of rank when compared to SAME clustering. CHAI-AvgSim and CHAI-SNF have a median rank of 3 and 2 respectively, compared to SAME clustering’s median rank of 5. Additionally, CHAI-AvgSim is the most consistent performer in terms of rank, with its lowest rank across datasets being 5, compared to CHAI-SNF’s lowest rank of 6 and SAME-Clustering’s lowest rank of 7.

### 2.3 CHAI outperforms existing clustering methods across varying dataset sizes and complexity

In order to evaluate CHAI on varying datasets in terms of complexity and size, we took advantage of the varying cell types and large number of cells in the Zheng68K PBMC dataset [26]. We created six different datasets, with three different sizes and number of cell types. We refer to the datasets with five equally sized groups as “simple” cases and randomly selected groups as “challenging”.

CHAI-AvgSim and CHAI-SNF are robust performers across dataset conditions, as seen in Fig. 3a. Both methods are top three performers in all six of the subsampled datasets; additionally, CHAI-AvgSim is the top performer in three of the six datasets. Either CHAI method has a better ARI than SAME-Clustering, the other consensus clustering method, in all six of the subsampled sets. In Fig. 3c, we note that CHAI-AvgSim has the highest median ARI, while CHAI-SNF has the lowest interquartile range. This suggests that CHAI-AvgSim calculates a higher ARI more frequently, but CHAI-SNF is more consistent in performance.

**Figure 3.**
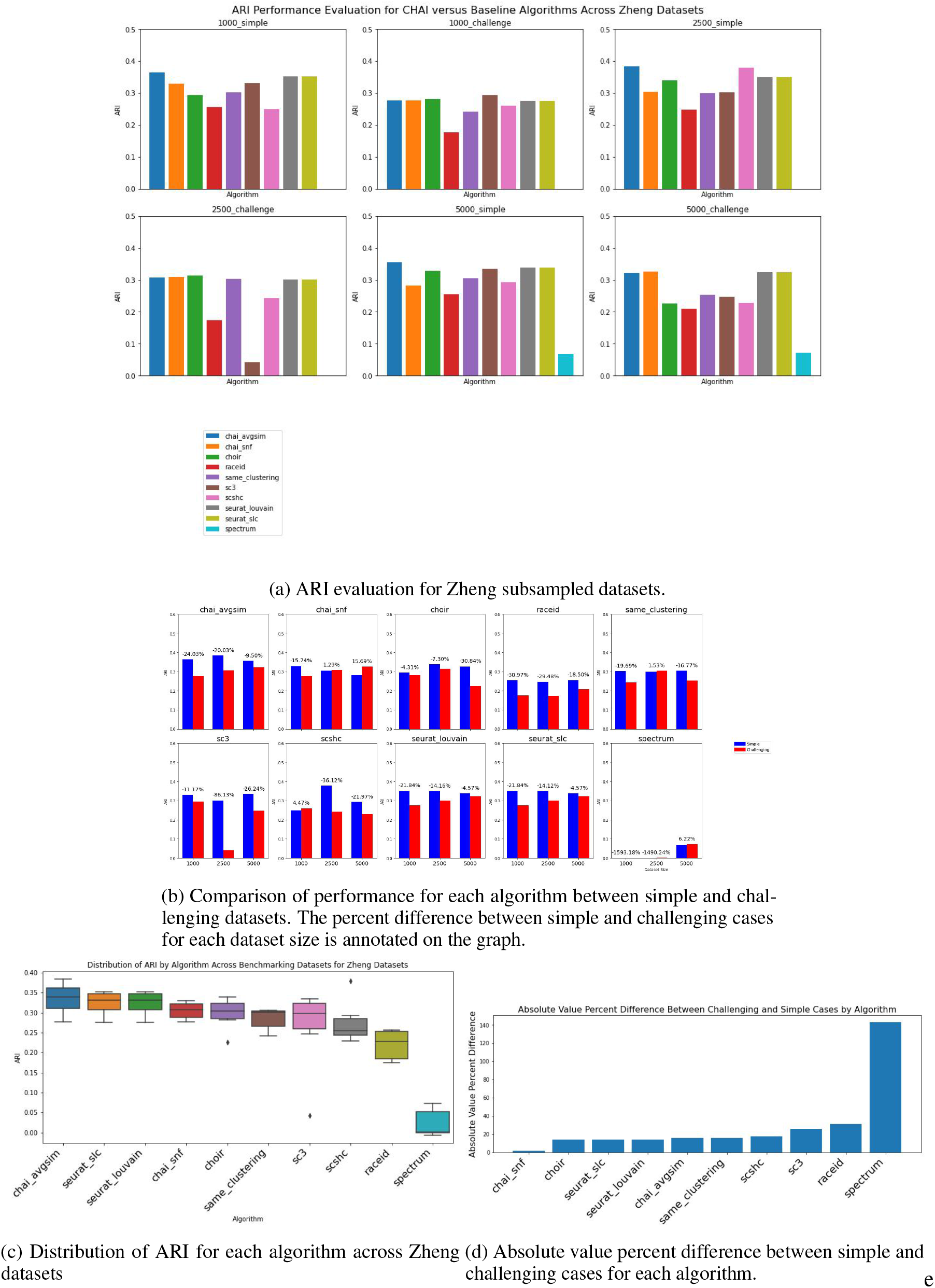
CHAI Evaluation on Zheng Subsampled Datasets

We also wanted to evaluate how well each method performs when faced with a simple or challenging dataset. Fig. 3b displays the percent difference between simple and challenging datasets for each algorithm across dataset sizes. Most algorithms decrease in performance in terms of ARI when evaluated on a dataset with randomly selected groups, across dataset size. Notably, CHAI-SNF seems to actually increase in performance on challenging datasets, even as the size of the dataset increases. We consider that a consistent algorithm would perform well when dataset sizes are the same, but the topologies of clusters are different. Therefore we examine the absolute value of percent difference across dataset sizes, but between the simple and challenging datasets, depicted in Fig. 3d. CHAI-SNF has very little difference between simple and challenging datasets; this is in contrast to CHAI-AvgSim, which has the highest median ARI and a low interquartile range, but displays a larger percent difference between its simple and challenging cases. Both methods ultimately outperform the other consensus method, SAME-Clustering, in terms of median ARI, consistent performance by ARI distribution, and low percent difference between simple and challenging cases.

### 2.4 CHAI derives validated biological insights in a breast cancer dataset: case study

A potential concern surrounding consensus clustering methods is that the features of certain methods may be over-shadowed by the results from all other methods. scRNAseq clustering methods use a variety of different techniques to determine the final cell to cluster assignments, which involve a varying degree of biological information [3]. Many methods, such as Seurat and CHOIR, filter the initial expression matrix through PCA and identify the highly variable genes within the dataset [4] [8]. Other methods, such as tSNE + KMeans Clustering, do not use any prederived biological insight prior to clustering [19]. There are also clustering methods, such as CIDER, which recluster cells based on differentially expressed gene (DEG) signature [27]. With this diversity in clustering in mind, we tested if CHAI can reliably derive biological conclusions as a standalone method. We decided to use CHAI-AvgSim for this analysis, as it demonstrated better consistency across dataset conditions than CHAI-SNF in our benchmarking.

Here we perform clustering on a dataset from Hwang et al., which studies collective cell migration of breast cancer [28]. During collective migration *in vivo*, breast cancer cells move as a cluster and prior work suggests that cells within the clusters can be heterogeneous [29]. Thus, Hwang et al. used single cell sequencing to identify different cell populations within collectively migrating clusters, with the ultimate goal to understand how cells at the front, known as leader cells, may have unique gene signatures that allow them to lead migration. To induce migration, Hwang et al used biochemical and biomechanical gradients, and performed single cell sequencing analysis after migration had occured (GEO Accession number: GSE171203) [28]. After induction of biochemical gradient stromal-derived factor 1 (SDF1), single cell sequencing analysis of tumor clusters revealed 9 different cell population types and 1 primary cluster of leader cells with differential expression of Cadherin-3 (CDH3) [28].

In our data validation, we analyzed the dataset for the cell clusters migrating in response to the biochemical gradient stromal derived factor 1 (SDF1), and refer to this dataset as “SDF1”. First, we performed consensus clustering using

CHAI-AvgSim on the SDF1 dataset, which also revealed 9 different clusters. To determine how accurately CHAI was able to identify leader cells in the SDF1 dataset, we compared percentage of shared cells between the ground truth clusters and the clusters predicted by CHAI-AvgSim. In Hwang et al’s single cell analysis, cluster 4 contained the leader cells, and we see in Fig. 4a that cluster 4 has greater than 90% cell overlap with CHAI-AvgSim Cluster 5. In other words, over 90% of the cells predicted to be in Cluster 5 from CHAI-AvgSim are in fact experimentally validated leader cells.

**Figure 4.**
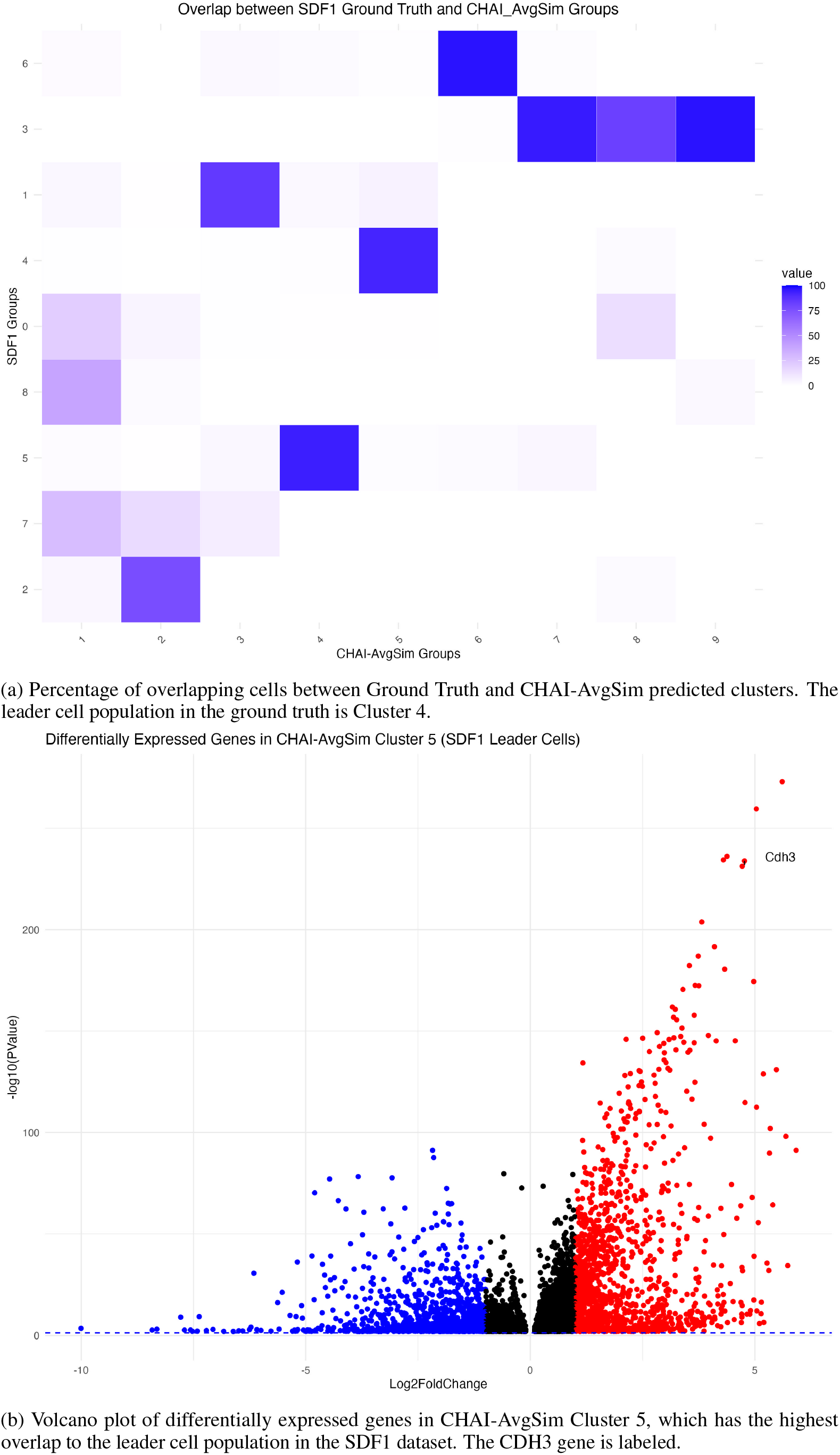
CHAI-AvgSim Analysis of CDH3 Leader Cell Population in SDF1 Induced Migration Dataset

**Figure 5.**
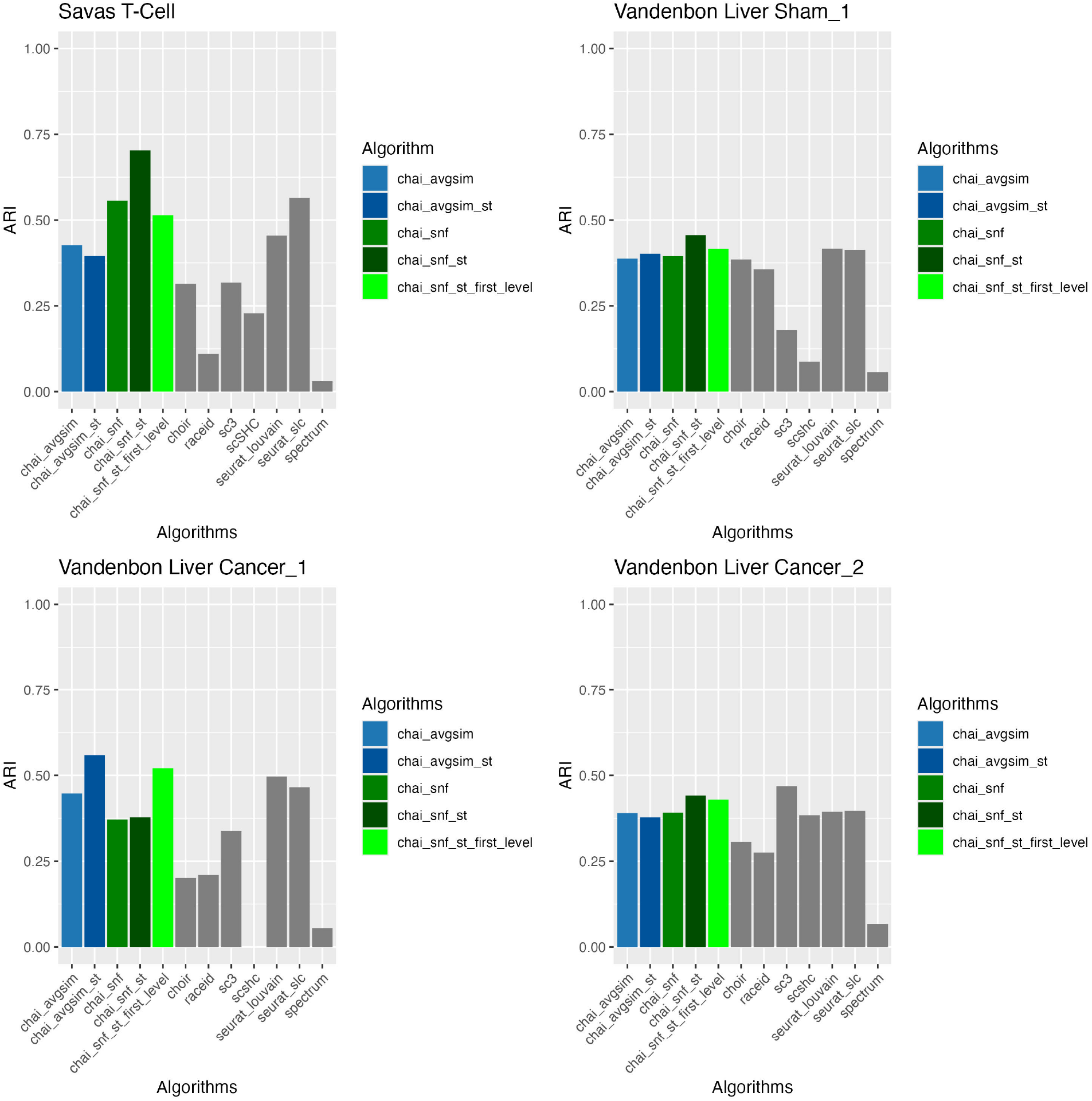
ARI Evaluation for CHAI Spatial Transcriptomic Integration. All CHAI spatial transcriptomics integrated are suffixed with “-st” in the bar labels.

To validate biological relevance of our approach, we calculated differentially expressed genes and visualized them using a volcano plot in Fig. 4b. We calculated the differentially expressed genes by running the FindAllMarkers function in Seurat [4]. The primary goal behind this analysis was to determine whether CHAI cluster 5 cell population was enriched for CDH3, a demonstrated leader cell marker in the original study [28], as a way to validate cluster 5 is indeed the leader cell population. Our analysis demonstrates CDH3 is significantly upregulated in the CHAI-AvgSim leader cell cluster, when compared to other clusters. Thus, CHAI-AvgSim was able to accurately identify the leader cell sub-population distinctly. This study demonstrates the accuracy of CHAI and validates its ability as a method to derive biological insights.

### 2.5 Integration of spatial transcriptomics data with CHAI: CHAI-ST

As CHAI relies on binary matrices to represent cell to cell relationships, we evaluated if other modalities may be integrated into the CHAI framework, provided that they can be represented as binary matrices. Spatial transcriptomics is an emerging sequencing technology that quantifies the location of a cell at the time of sequencing [30]. A recently published method, GraphST, is able to represent the relationship between cells based on their spatial coordinate distance as a binary matrix [22]. We extend this approach from GraphST and easily integrate it into the proposed CHAI framework. The main purpose of this experiment was to quantify if the incorporation of other data modalities to CHAI will improve the overall clustering accuracy.

We present several options to integrate spatial transcriptomics into the CHAI package. For CHAI-AvgSim, we integrated the spatial transcriptomics data by simply including it in the average matrix calculation as another modality. For CHAI-SNF, we first ran SNF on the clustering algorithm binary matrices. We then ran SNF again on the clustering algorithm SNF matrix and the binary matrix from the spatial transcriptomics data, therefore running two levels of SNF. Finally we run CHAI-SNF-First-Level, in which we incorporate the spatial transcriptomics binary matrix alongside the binary matrices of the other clustering algorithms, and run SNF just once to determine the final clustering assignments [20].

We evaluated CHAI with the integration of spatial transcriptomics coordinates on four datasets using ARI. From this analysis, we find that the integration of spatial transcriptomics with CHAI-SNF improves the ARI in all four datasets. Additionally, we see that the integration of spatial transcriptomics causes either CHAI method to be the top performing algorithm in three out of the four datasets. The ARI for CHAI-AvgSim stays relatively the same when including spatial transcriptomics in most datasets, except for the Vandenbom Liver Cancer dataset, where the integration of the additional data significantly aids its performance. From this analysis we conclude that it is best to include spatial transcriptomics with CHAI-SNF. We see that incorporating the spatial coordinates separately and running two levels of SNF leads to better ARI in three of the four datasets. There is also no downside to including spatial transcriptomics data with CHAI-SNF or CHAI-AvgSim if available; even if the results do not significantly improve, we see that adding the additional information will still keep the ARI approximately the same.

## 3 Discussion

Clustering for scRNAseq data is a common task that has a variety of approaches. Each method has their own individual strengths and weaknesses, and there is currently no one best method that works with definitive superiority in all situations. This conclusion has been drawn from several benchmarking studies, including the one we put forward in this study [3]. Other ensemble clustering methods have been applied for scRNAseq data, but these are based on older versions of scRNAseq clustering methods and have not been updated or maintained frequently [31] [19]. With CHAI-AvgSim and CHAI-SNF, we present two distinct consensus clustering methods that each have their own advantages. Both methods demonstrate improved performance on several dataset conditions and complexities.

First, we chose 10 benchmarking datasets to evaluate both CHAI-AvgSim and CHAI-SNF on and compared them with the individual clustering algorithms that made up the consensus pipeline. We found that CHAI-SNF has the highest median ARI across all of the dataset runs, and the highest maximum ARI as well. However CHAI-AvgSim demonstrates comparable median ARI, while also having the lowest interquartile range out of all of the other algorithms. This, combined with the fact that CHAI-AvgSim is a top three performer in 80% of all benchmarking datasets, suggests that it is a more consistent and safer choice to use when the exact structure of a dataset is not known. We note the variation across all of the datasets in most of the algorithms. The previous consensus clustering method we chose to compare to, SAME clustering, has a similar median ARI and interquartile range when compared to both CHAI-AvgSim and CHAI-SNF. However it has a much lower median rank, and does not feature as regularly in the list of top 3 performers across datasets. When evaluated on simple and challenging cases, both CHAI-AvgSim and CHAI-SNF show consistency between the two cases. We note that CHAI-SNF has a significant percent difference between its simple and challenging cases, across all dataset sizes. From this analysis, we are able to conclude that CHAI-SNF is least susceptible to varying performance as dataset complexities increase.

When comparing both CHAI methodologies to SAME-Clustering, it is important to note that we used the current version of SAME-Clustering available, where SC3 does not run in its package due to a bug (see: https://github.com/yycunc/SAMEclustering/issues/4. Therefore SC3 is included in our pipeline, while not being included in SAME-Clustering’s in all of the evaluations we conducted [5] [19]. Despite this fact, we are still confident of CHAI’s performance as it incorporates several other algorithms that are not included in SAME-Clustering. Users may also notice Spectrum’s poor performance, often displaying subzero and negative ARI [6]. We included Spectrum anyways to demonstrate that CHAI’s performance is overall unaffected by a singular poor performing algorithm, provided that the rest of the algorithms demonstrate a reasonable accuracy. As more clustering algorithms are added and the community continues to see variable performances, CHAI will remain to be a stable choice unlikely to be influenced by one singular extremely poor performing algorithm.

When gold standard cell types are not available, we sought to demonstrate CHAI’s practical usability for identifying important clusters and biomarkers in a real world application. We found that CHAI was able to identify a CDH3 enriched cell population which has been linked to leading cell migration in breast cancer [28]. This demonstrates that not only does CHAI have a better performance in terms of accuracy, it is also able to derive biologically meaningful results.

As multiomic data for single cell genomics increases, the need to integrate this information will continue to arise [32]. In this study, we choose spatial transcriptomic coordinate data as an example for multiomic integration with CHAI. Using a binary similarity matrix method developed from GraphST, we show that adding this additional omic to CHAI-AvgSim increases it significantly in one benchmarking dataset, and keeps performance relatively the same in the other datasets [22]. For CHAI-SNF on the other hand, the integration of spatial transcriptomic data increases the performance in all cases. As the original purpose of SNF was to integrate disparate modes of data for the same sample, this makes CHAI-SNF a logical choice for this purpose [20]. The nature of CHAI allows for it to accommodate other forms of data, so long as they can be represented as a binary similarity matrix between cells. This makes it a generalized method for not only standard clustering, but multiomic clustering as well. The flexibility of the binary matrix architecture will lend CHAI usable in a variety of different purposes going forward.

Further evaluation remains to be done on the best algorithms to use in the consensus pipeline for a particular dataset conditions. As more methods emerge and quality benchmarking is performed, various sets of algorithms will become the best performers given a specific dataset. In these instances we aim for CHAI to be customizable, where several algorithms can be added or removed based on user preference. Ideally these choices will be informed by community best practices. However based on current evaluations, it is our recommendation to include as many algorithms as possible.

## 4 Conclusion

We present CHAI, a consensus clustering method demonstrating robust and superior performance in a wide variety of dataset conditions for scRNA-seq data. CHAI is able to detect key biomarkers in cancer tumor cells; additionally, CHAI provides a platform for multiomic integration. We hope CHAI is a tool for the community, where new algorithms may be integrated seamlessly and other omics are built into the pipeline.

## 5 Methods

The CHAI workflow may be summarized as three majors steps:

1. Run individual clustering algorithms and compute binary similarity matrix for each.
2. Calculate Average Similarity matrix and/or SNF matrix
3. Run Spectral Clustering on either integrated matrix to determine final cell identities.

The package is written in R, and is available for installation on GitHub at https://github.com/lodimk2/chai.

### 5.1 Individual Clustering Algorithms

CHAI incorporates seven algorithms by default when using the package, which are described below. Users may also integrate information from other clustering methods.

#### 5.1.1 Seurat

Seurat begins with reducing the dimension using methods such as PCA, UMAP, and tSNE. It then identifies variably expressed genes, then a K nearest neighbor (KNN) graph is computed based upon these. From here, community detection algorithms are used to identify the final clusters. Both Louvain and SLC rely on the local moving heuristic for modularity optimization. The premise is to continually move individual nodes from one community to another so that each node movement elicits a modularity increase. This is done in a random order. For each node, it is checked whether it is possible to increase the modularity by moving it to a different community. If this is possible, then the node is moved to the community that results in the highest modularity gain. This repeats until it is no longer possible to increase modularity through individual node movements. In CHAI, we used Louvain and SLC. There are two versions of Louvain that are used in the paper: Louvain and Louvain with Multilevel Refinement. Both algorithms follow the same steps, with the difference being that the local moving heuristic is run again at the end of the program to fine-tune the final community structure and to also guarantee that the final community structure can not be further optimized. First, an adjacency matrix of a network and the initial assignments of nodes to communities is inputted. The local moving heuristic is run. If the number of communities is less than the number of nodes, then a reduced network is created. A recursive call is then performed to identify the community structure of the reduced network. The communities are then merged based off this community structure. Finally, based off which version of Louvain is run, the local moving heuristic can be performed. SLC applies the local moving heuristic differently than Louvain. First, the local moving heuristic is run. Then, if the number of communities is less than the number of nodes, a subnetwork for each community is created and the local moving heuristic is run for each subnetwork. A reduced network is then formed based on the community structure of the subnetworks. A recursive call is performed to identify the community structure of the reduced network, and the communities are merged based on those findings.

#### 5.1.2 CHOIR

CHOIR constructs a hierarchical clustering tree. Using all cells, it identifies a set of features that have variable levels of expression. Then, dimensionality reduction is applied using either PCA, LSI, or iterative LSI, with PCA being the default method. A nearest neighbor adjacency matrix is computed, and to generate the layers of the clustering tree, Louvain and Leiden clustering is used. MRtree is used to reshape the clustering trees into a hierarchical tree [8].

#### 5.1.3 RaceID

RaceID uses K-means clustering. First, a similarity matrix is constructed, which contains Pearson’s correlation coefficients for all pairs of cells. K-means clustering is then applied to it, and the number of clusters used for k-means clustering is decided on by the difference of the average within cluster dispersion in the data. It also computes Jaccard’s similarity to check if fewer clusters should have been produced [21].

#### 5.1.4 SC3

SC3 uses a gene filter to remove any genes or transcripts that are in less than X% of cells (X being commonly set to 6). After calculating the distance between the cells, using Euclidean, Pearson and Spearman metrics, all distance matrices are then transformed. K-means clustering is then applied. A consensus matrix is computed using CSPA (Cluster-based Similarity Partitioning). For each individual cluster result, a binary similarity matrix is made. If two cells belong to one cluster, their similarity is 1; otherwise, it is 0. The consensus matrix is created by averaging all similarity matrices of the individual clusterings. [5].

#### 5.1.5 Spectrum

Spectrum uses an adaptive density-aware kernel (based on the Zelnik-Manor self-tuning kernel and the Zhang density-aware kernel) to construct the similarity matrices. These matrices are combined using TPG diffusion. Then, the Ng spectral clustering method is applied to the similarity matrix [6].

#### 5.1.6 scSHC

scSHC used hierarchical clustering as a part of their algorithm. The first step is to compute the distance between each cell, but since scRNA-seq data has small counts and high dimensionality, finding the Euclidean distance is unreliable. Therefore, Euclidean distance on the latent variables is computed instead. To identify the clusters, a desired family-wise error rate is decided upon (0.05 in simulated data and 0.25 on real data applications). The method goes down the tree to decide which splits should be kept. This decision is made using hypothesis testing: a test statistic is formed using the average silhouette, which is then compared to the desired family-wise error rate. If it is greater or equal to the desired family-wise error rate, then it failed to reject the null hypothesis, and all data should belong to one cluster. Otherwise, the data is split into the two proposed clusters and the method continues down the tree [9].

### 5.2 CHAI-AvgSim

Once the individual clustering assignment algorithms are run, they will each be represented as a table containing the Cell ID in one column, and the Clustering Assignment as the other column. From here, we convert this table to a binary similarity matrix. We represent a cell to clustering assignment vector as a binary similarity matrix using the following rules:

1. If two cells have the same clustering assignment, assign a value of 1 to a binary similarity matrix corresponding to the two cells.
2. If two cells do not have the same clustering assignment, assign a value of 0 to the binary similarity matrix corresponding to the two cells.

Through this method, each clustering assignment is converted into a binary similarity matrix. Each binary similarity matrix per algorithm is then aggregated into an Average Similarity matrix, which simply put is a cell to cell correlation matrix containing the per element average rank across all individual clustering algorithm matrices.

Consider a dataset with *m* cells. Therefore, each binary similarity matrix per algorithm will be of dimension *m× m*. To construct an Average Similarity matrix of *m×m* dimension, we calculate the average per cell using the following formula:

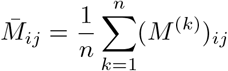

Where:

*n* = Total number of samples

*n*_*ij*_ = Number of pairs of samples that are in the same cluster in both the first and second clusterings

*a*_*i*_ = Number of samples in the *i*th cluster according to the first clustering

*b*_*j*_ = Number of samples in the *j*th cluster according to the second clustering

This formula is repeated across each cell in the matrix until a final *m × m* matrix is created.

Once the Average Similarity matrix is computed, we use Spectral Clustering to determine the final cell clusters [33]. If the true number of clusters is known to the user, they can use this *k* value as the number of partitions to make on the Average Similarity matrix. If the true number of *k* in the dataset is not known to the user, we recommend calculating the *k* value for which the silhouette score is the highest. For all evaluations conducted in our benchmarking, we conduct a silhouette score evaluation in range 2 to *k* + 1, with *k* being the true number of clusters present in the dataset. Despite the true number of clusters being known in the benchmarking dataset, we choose a value of *k* computationally in order to simulate working with unknown data.

### 5.3 sCHAI-SNF

The CHAI-SNF method begins similarly to CHAI-AvgSim, where a clustering table containing Cell ID and Clustering Assignment is converted into a binary similarity matrix for each clustering algorithm. However, rather than taking an average vote across cell to assignment similarities, we apply the Similarity Network Fusion (SNF) algorithm across all binary similarity matrices [20].

The SNF algorithm was created for multiomic data integration in bulk RNA sequencing data. It was used to integrate patient to patient similarity matrices across three data modalities: mRNA expression, DNA methylation and microRNA (miRNA) expression. Once the matrices were integrated, the final matrix was used for downstream tasks such as cancer subtyping and survival analysis [20]. In brief, SNF performs similarity matrix fusion by converting a pairwise patient similarity matrix to a graph, where nodes are the patients and edges are the relationships between the patients. From here SNF uses a network fusion step based on message passing theory that iteratively updates each network, which makes it more similar to the other networks until all networks are the same. SNF has been demonstrated to remove low edge weights, also known as “weak edges”, from the final network, and include only relationships that are more likely to be in concordance with the ground-truth [20].

Ultimately, since we have cell to cell similarity matrices for each clustering algorithm, applying SNF to the individual algorithm’s binary similarity matrix representation was straightforward. We implemented SNF using the SNFtool package in R, available on CRAN, using the default parameters. For more detailed information on SNF, please refer to Wang et al. [20].

Similar to CHAI-AvgSim, we infer the final clusters by running Spectral Clustering on the final SNF combined matrix, either by knowing the true *k* value or by calculating the best *k* by silhouette score optimization.

### 5.4 GraphST Binary Matrix Representation for Spatial Transcriptomics

GraphST is a method that integrates spatial coordinates with scRNAseq data. One step in their process is to represent the distance between cells as a binary matrix [22]. We incorporate that logic here into CHAI in order to integrate spatial transcriptomics into CHAI-AvgSim and CHAI-SNF.

GraphST creates an undirected neighborhood graph represented as a binary adjacency matrix, where the number of neighbors to any one cell is set to be a predefined number *k*. The neighbors of a spot *s* ∈ *S*, where each spot is represented as a vertex of the graph, represent the *k* spatially closest spots to *s*. Enumerating *S*, the adjacency matrix *M* ∈ ℝ^*n×n*^, where *n* is the number of spots, is constructed such that *a*_*ij*_ = 1 if *i, j* ∈ *S* are neighbors and 0 otherwise [22].

A neighborhood matrix created utilizing the same logic is incorporated into CHAI-AvgSim as another clustering assignment in the average matrix. Additionally, after applying CHAI-SNF on the various clustering assignments to produce a preliminary clustering assignment matrix, SNF is applied once again on this resultant matrix and the created neighborhood matrix to obtain the final clustering matrix that incorporates spatial data.

### 5.5 Evaluation Metrics

#### 5.5.1 Adjusted Rand Index (ARI)

Adjusted Rand Index (ARI) is a frequently used evaluation metric for clustering data, particularly in single cell genomics clustering [31]. ARI measures the concordance between a predicted set of clusters and the true set of clusters, scaled between -1 and 1. The higher the ARI, the better the performance, with 1 indicating a perfect overlap between the predicted and true clusters [34].

ARI may be calculated using the following formula:

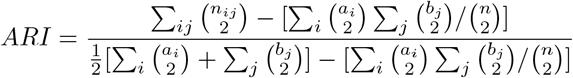

Where:

*n* = Total number of samples

*n*_*ij*_ = Number of pairs of samples that are in the same cluster in both the first and second clusterings

*a*_*i*_ = Number of samples in the *i*th cluster according to the first clustering

*b*_*j*_ = Number of samples in the *j*th cluster according to the second clustering

#### 5.5.2 Normalized Mutual Information (NMI)

Normalized Mutual Information (NMI) is a measure used to quantify the similarity between predicted clusters and the true clusters. It stems from the concept of mutual information, which measures the amount of information obtained about one random variable through the observation of another random variable. NMI ranges from 0 to 1, where 0 indicates no mutual information between the predicted and true clusters, and 1 indicates perfect agreement between the predicted and true clusters [35].

The mutual information between the predicted and true clusters, *C* and *K* is given by:

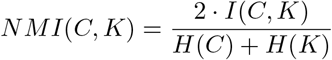

Where:

*NMI*(*C, K*) = Normalized Mutual Information between clusterings *C* and *K*

*I*(*C, K*) = Mutual Information between clusterings *C* and *K*

*H*(*C*) = Entropy of clustering *C*

*H*(*K*) = Entropy of clustering *K*

#### 5.5.3 Silhouette Score

To evaluate the best *k* for Spectral Clustering on either the CHAI-AvgSim or CHAI-SNF matrix, we calculate the best average Silhouette Score. Silhouette score measures how close each sample in one cluster is to the samples in neighboring clusters, which helps to assess the quality of clustering. This metric ranges from -1 to 1, with a high score indicating a cell is matched closely to its labeled cluster. Silhouette Score is calculated using the following formula:

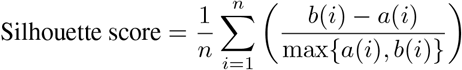

Where:

*n* is the total number of samples

*a*(*i*) is the average distance from sample *i* to other points in the same cluster

*b*(*i*) is the smallest average distance from sample *i* to points in a different cluster

## 6 Data

### 6.1 Baron Pancreas Data

Baron et al. addresses the limitations of previous gene expression profiling in the pancreas by using a droplet-based, single-cell RNA sequencing method to analyze over 12,000 individual pancreatic cells from 4 human donors and 2 mouse strains [36]. The analysis demonstrated 15 distinct clusters of cells, including subpopulations which were validated through immunohistochemistry. Additionally, heterogeneity was observed within human beta-cells, highlighting differences in gene regulation related to functional maturation and endoplasmic reticulum stress. Leveraging single-cell data, the researchers detected disease-associated differential expression and identified novel cell type-specific transcription factors and signaling receptors [36]. Over the years, the Baron dataset has served as a resource for validating and comparing findings in single-cell RNA sequencing studies because it is a large dataset with a view of gene expression patterns across distinct cell types [37]. You may download the data through GEO with accession number GSE84133.

### 6.2 Muraro Pancreas Data

Few proteins uniquely distinguish cells within the pancreas, creating a challenge because traditional techniques like immunohistochemistry rely on specific markers and may not sufficiently distinguish various cell populations. Muraro et al. describes using an automated platform that combines Fluorescence-Activated Cell Sorting (FACS), robotics, and the CEL-Seq2 sequencing protocol [38]. This approach allowed them to obtain transcriptomes from thousands of single pancreatic cells from deceased organ donors. As a result, they were able to identify cell type-specific transcription factors, discover a subpopulation of REG3A-positive acinar cells, and establish CD24 and TM4SF4 as markers for sorting alpha and beta cells. (GEO accession number: GSE85241)

### 6.3 SC-Mixology Data

The SC-Mixology dataset involves three human lung adenocarcinoma cell lines: HCC827, H1975, and H2228. Single cells from each cell line were processed using CEL-seq2, Drop-seq, and 10X Chromium library preparation methods then sorted into 384-well plates. Additionally, bulk RNA from each cell line was mixed in different ratios, diluted to single-cell equivalents, and sequenced [25]. The data is downloadable from the authors’ Github: https://github.com/LuyiTian/sc_mixology

### 6.4 Zeisel Mouse Brain

Zeisel et al. utilized single-cell RNA sequencing to analyze 3,436 mouse brain and 1,504 lung cell transcriptomes, aiming to understand vascular diseases. They identified 15 distinct cell clusters in the brain cortex and hippocampus and 17 in the lung, providing insight on tissue cellular diversity and organization [39] (GEO accession number: GSE103840)

### 6.5 Zheng 68K PBMC Data

The Zheng68K dataset by 10X CHROMIUM is a large dataset consisting of 68,450 blood mononuclear cells. The dataset was developed using an adaption of GemCode single-cell technology. There are eleven subtypes of cells within this dataset, those being CD8+ cytotoxic T cells (30.3%), CD8+/CD45RA+ naive cytotoxic cells (24.3%), CD56+ NK cells (12.8%), CD4+/CD25 T Reg cells (9.0%), CD19+ B cells (8.6%), CD4+/CD45RO+ memory cells (4.5%), CD14+ monocyte cells (4.2%), dendritic cells (3.1%), CD4+/CD45RA+/CD25-naive T cells (2.7%), CD34+ cells (0.4%) and CD4+ T Helper2 cells (0.1%). For CHAI benchmarking, we took advantage of the diversity contained in the Zheng68K dataset by subsampling it into six smaller datasets, those being:

1. 1000 cells with 5 equal populations
2. 1000 cells with random populations
3. 2500 cells with 5 equal populations
4. 2500 cells with random populations
5. 5000 cells with 5 equal populations
6. 5000 cells with random populations

From this subsampling analysis we were able to benchmark CHAI against varying dataset conditions and controls [26]. We consider the datasets with equal populations to be “simple” datasets, and with random groups to be “challening” datasets.

### 6.6 Savas Breast Cancer T Cell Data

Savas, P. et al [40] studied the characteristics of T cells in breast cancer tumor-infiltrating lymphocytes (TILs). Multi-parameter flow cytometry was utilized to analyze breast cancers for their TIL content. Data was obtained from 84 individuals with primary breast cancers and 45 individuals with metastatic breast cancers. The findings revealed significant heterogeneity in the infiltrating T cell population and suggested that CD8+ tissue resident memory T (TRM) cells contribute to breast cancer immunosurveillance and are primarily modulated by immune checkpoint inhibition.

The dataset used in this paper was obtained by performing single cell RNA sequencing on 5759 purified CD3+ single T cells passing quality control from two primary triple negative breast cancer (TNBC) patients, encompassing a total of 15623 genes and 11 different gene expression annotations. The spatial coordinates of the cells obtained from the tissue are also recorded. Data used can be downloaded from Broad Institute’s Single Cell Portal with accession number SCP2331.

### 6.7 Vandenbon Mouse Liver Cancer Visium Data

Zonation refers to the spatial organization of gene expression within the liver such that hepatocyte functions are specified by relative distance to the bloodstream. In [41], Vandenbon et. al utilized spatial transcriptomics in order to investigate the quantity and zonation of hepatic genes in mice with cancer with the intention of determining whether liver zonation is influenced by solid cancers. This study found that liver zonation was influenced by breast cancers, exemplified by affected xenobiotic catabolic process genes, zonally elicited acute phase response, and zonally activated innate immune cells in the liver. Breast cancers zonally influencing liver gene expression profiles results in zonal liver functions also being affected. Data for this study was obtained from wild-type female mice. Four mouse liver samples consisting of two 4T1 cancer-bearing mice samples, Cancer1 and Cancer2, and two sham samples, Sham1 and Sham2, were processed with 10x Genomics Visium spatial transcriptomics, culminating in a dataset with a total of 7,758 spots and 32,285 genes clustered into 13 cell type categories.

For this case study, the Cancer1 (2110 spots), Cancer2 (1438 spots), and Sham1 (1952 spots) samples were utilized. The data used can be downloaded from Broad Institute’s Single Cell Portal with accession number SCP2046.

## 7 Code Availability

CHAI is available as an R package here: https://github.com/lodimk2/chai

