## Supplementary Figures for "CHAI: Consensus Clustering Through Similarity Matrix Integration for Cell-Type Identification"

March 19, 2024

### 1 Supplementary Figures

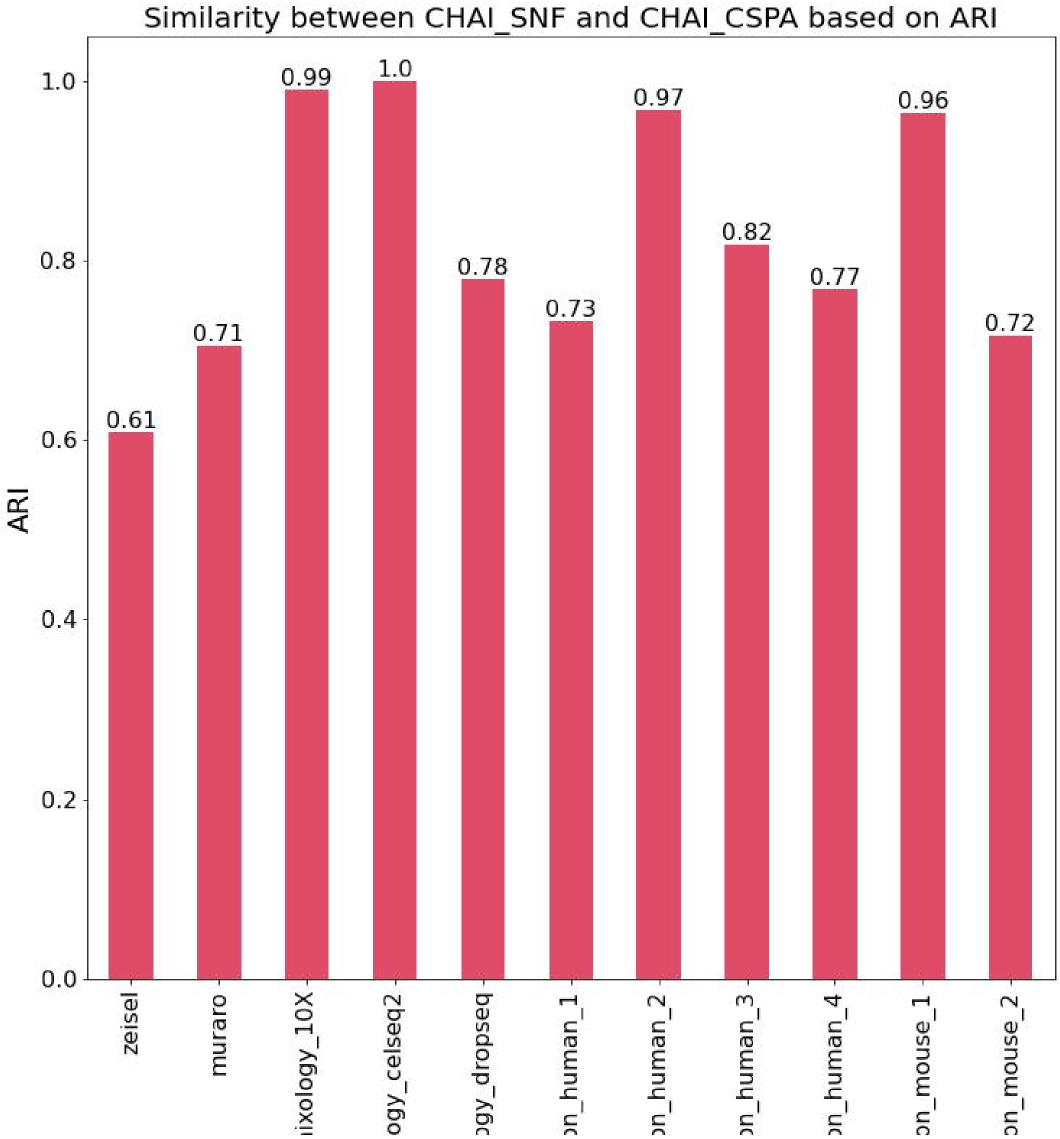


Figure S1: ARI Comparison between CHAI-SNF and CHAI-AvgSim by ARI. Despite differences in the ground truth evaluation, the similarity between clustering assignments by CHAI-AvgSim and CHAI-SNF is quite similar

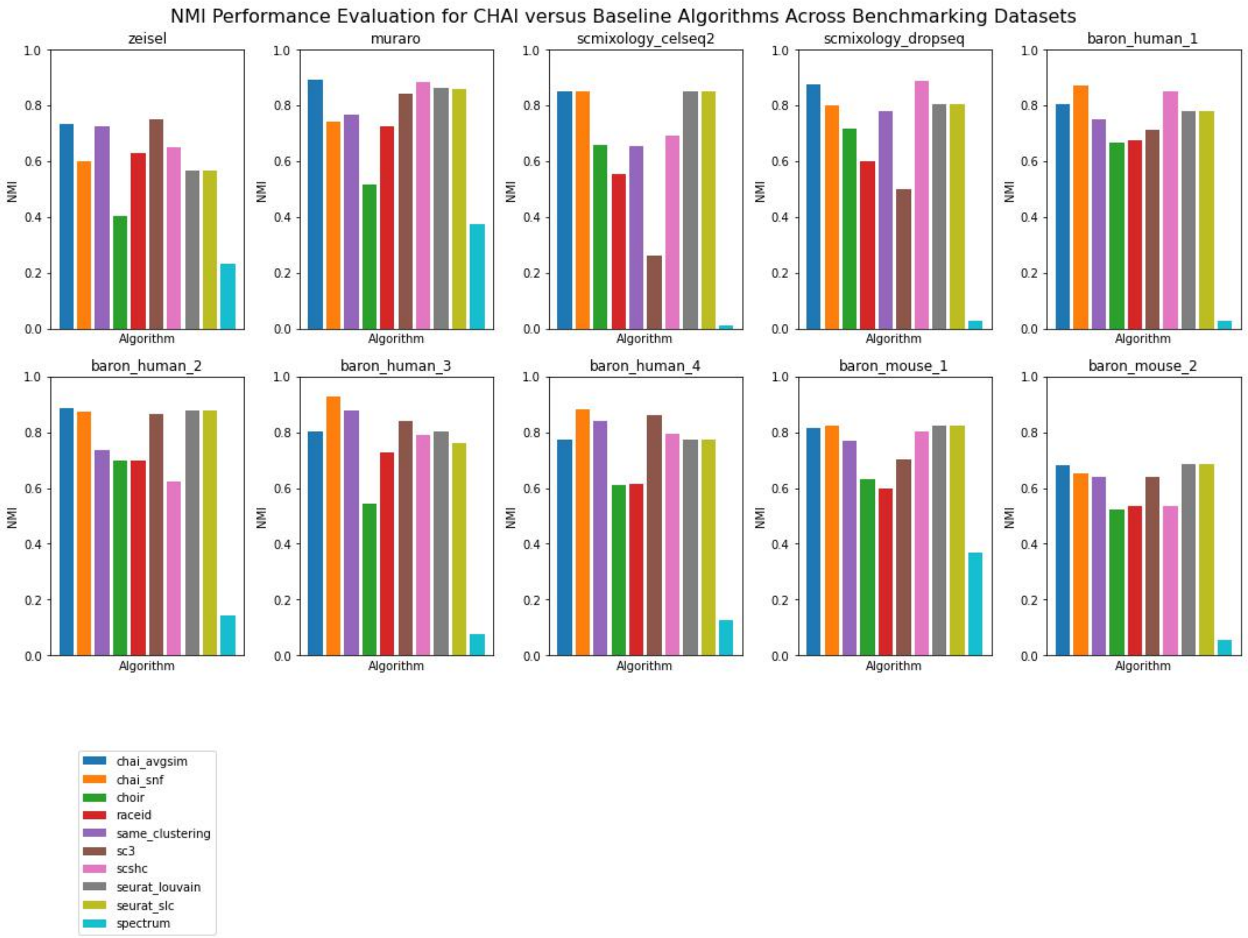


Figure S2: NMI Evaluation for Benchmarking Datasets

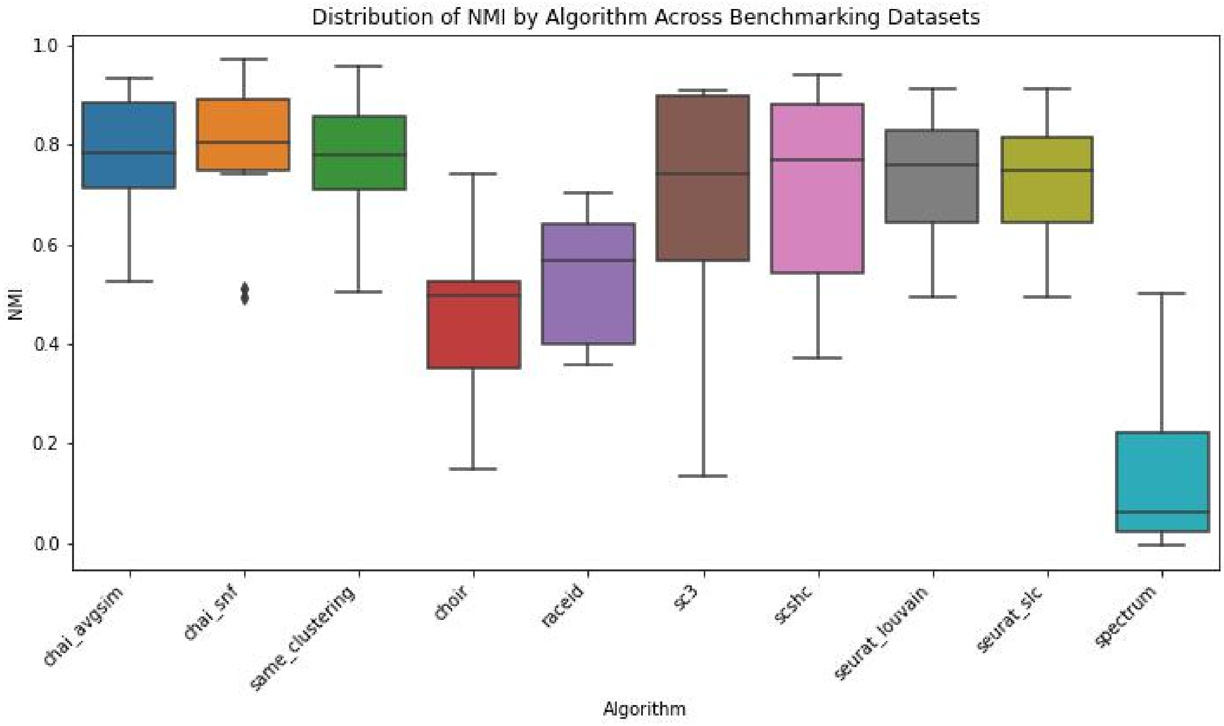


Figure S3: NMI Distribution for Benchmarking Datasets

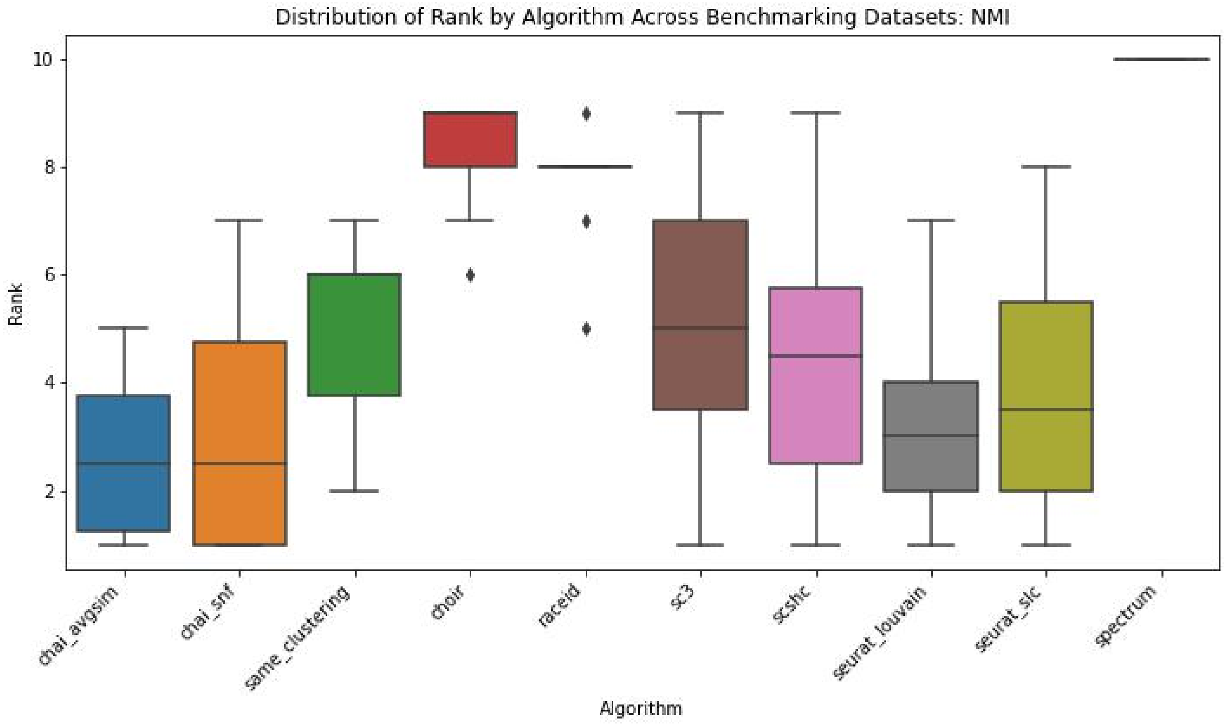


Figure S4: NMI Based Ranking Distribution for Benchmarking Datasets

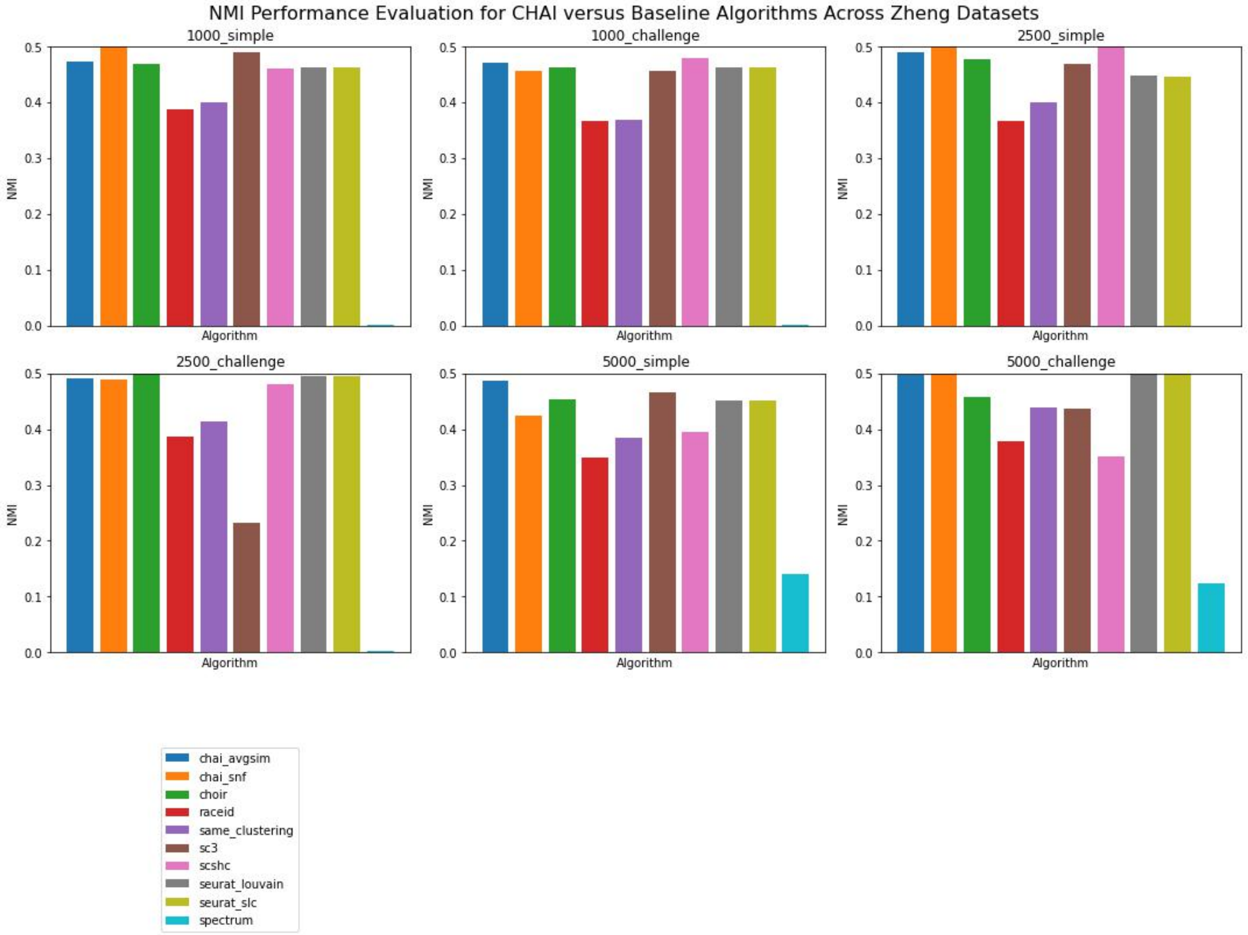


Figure S5: NMI Evaluation for Zheng Datasets

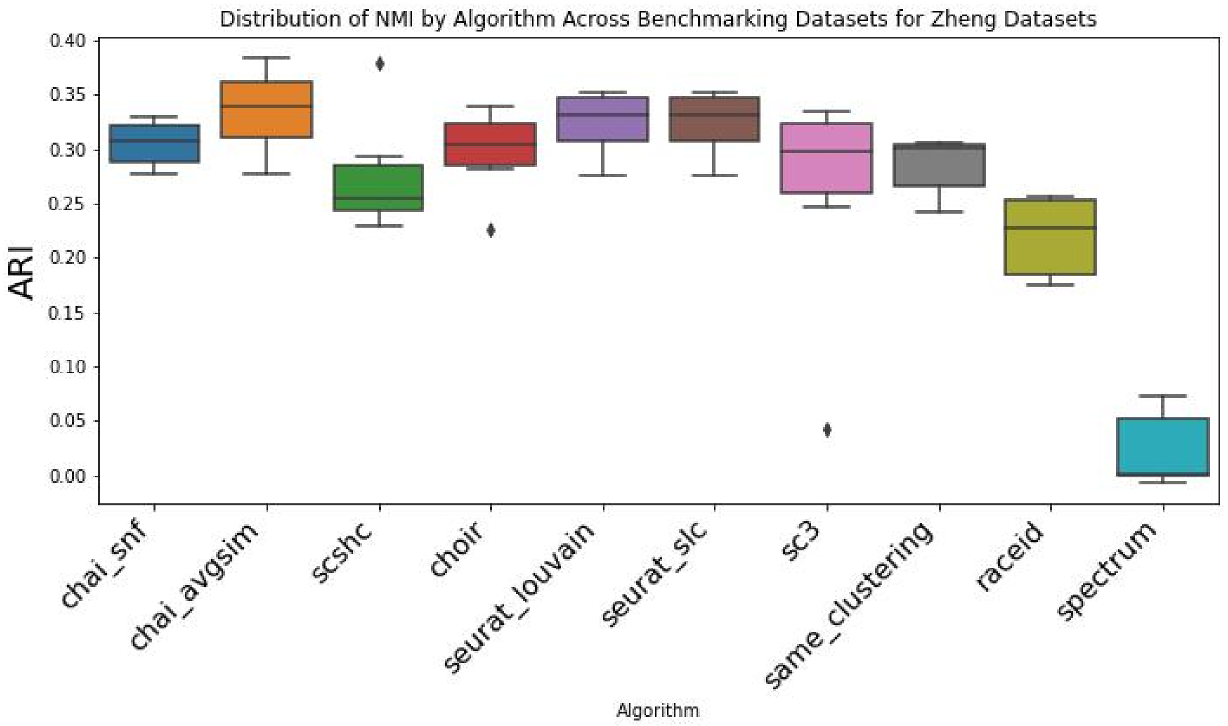


Figure S6: NMI Distribution for Zheng Datasets

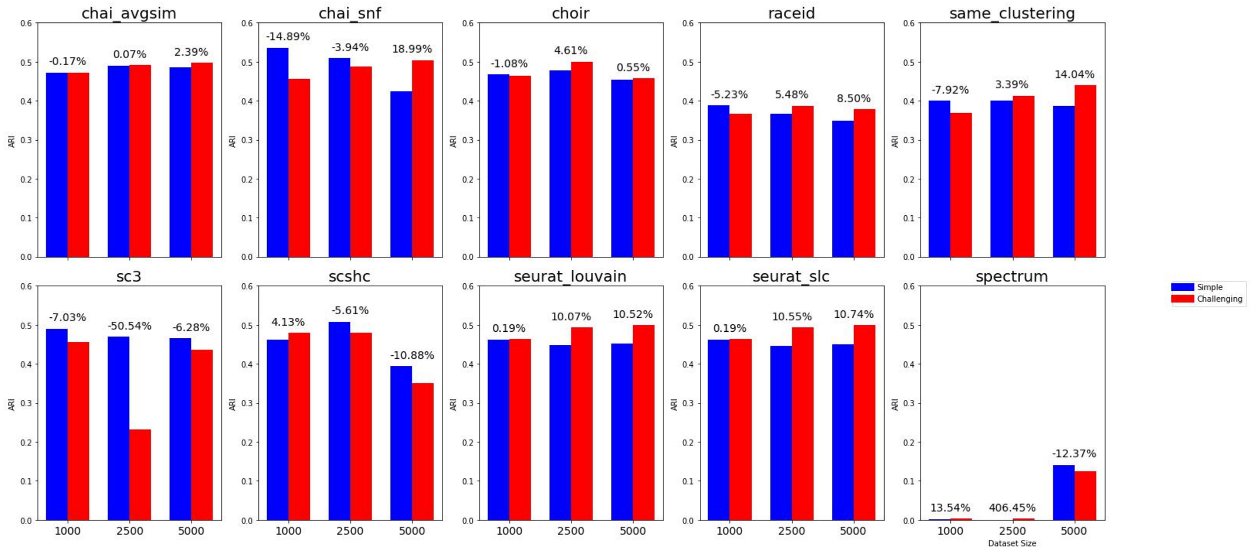


Figure S7: NMI Difference Between Simple and Challenging Datasets for Zheng Datasets
